# Predicting Post-Stroke Aphasia Speech Performance from Multimodal Data with Explainable Machine Learning

**DOI:** 10.64898/2026.02.02.703416

**Authors:** Shreya Parchure, Arnav Gupta, Apoorva Kelkar, Leslie Vnenchak, Olufunsho Faseyitan, John D. Medaglia, Denise Y. Harvey, H. Branch Coslett, Roy H. Hamilton

## Abstract

Aphasia, an acquired language deficit, is the most common post-stroke focal cognitive impairment, and roughly 60% cases become chronic (duration >6 months). Aphasia therapies could be optimized if clinicians could make personalized predictions of how individual persons with aphasia (PWA) would be likely to perform on particular language tasks. However, current approaches relying on imaging, lesion volume, patient demographics, and clinical scores achieve less than 50% accuracy in predicting performance in PWA. Research algorithms using complex imaging and fMRI can make binary predictions about the presence or absence of aphasia but do not give more clinically relevant information. We aim to predict word-by-word speech accuracy in PWA to better enable personalized speech therapies. To be clinically informative, machine learning models developed for this purpose should use clinically available inputs, explain key features behind a prediction, and generalize to new PWA and previously unseen words.

This study combines multimodal input features from clinical testing scores and structural MRI neuroimaging with a novel data source: word-by-word linguistic difficulty. We computed metrics of cognitive burden, such as semantic selection and recall demands, and articulatory burden, such as word length in phonemes and syllables, using naturalistic corpora containing over a billion words of English text. Retrospective training, ten-fold cross validation and 500-run bootstrapping of different machine learning models with various combinations of input features was conducted using 4620 trials. A simplified version of the best model using widely available inputs was deployed clinically through a web app, and prospective generalization was tested on 570 trials with unseen words and different naming tasks in new PWA.

We found the best performances with random forest classifiers using linguistic difficulty combined with either clinical information (AUROC ± SEM = 0.87 ± 0.07), or all together with structural imaging connectivity (0.90 ± 0.04). Classifiers using multimodal inputs significantly outperformed others employing single inputs (range 0.66-0.85, p<0.05). Extracting feature importances from the best model showed that Western Aphasia Battery scores, semantic demands, number of phonemes, and syllables were predictive of PWA speech accuracy. Structural integrity in peri-lesional brain regions predicted better language performance whereas higher connectivity of select contralateral homotopes contributed to prediction of worse speech. Without the inclusion of MRI data, lesion volume was a key predictor of PWA speech as well.

A simplified, clinically ready, explainable model (publicly available as AphasiaLENS web application) predicted PWA accuracy for any user-entered word, not restricted to a standardized battery. Its prospective generalization performance was not significantly different from the best model using full inputs (AUROC ranges 0.81-0.89, p>0.05). Thus, our research can help inform individualized treatment planning for PWA, while also suggesting research targets through better understanding of brain-behavior relationships.

## Introduction

Aphasia, or acquired impairment in language function, is the most common focal cognitive deficit after a stroke.^1–4^ Chronic aphasia, defined by deficits that persist beyond 6 months after a stroke, occurs in up to two-thirds of persons with initial acute post-stroke aphasia.^5^ This condition often greatly reduces quality of life and increases healthcare costs more than stroke alone^6^ because it disrupts individuals’ functional independence, social interactions, employment opportunities, emotional well-being, and healthcare delivery.^7–10^ Recent work suggests that persons with chronic aphasia can recover with personalized speech therapy^11–13^. An ideally individualized therapy would be one in which the ability of a person with aphasia (PWA) to correctly negotiate any given word could be known ahead of time, in order to best tailor treatment to that person. However, current neurological assessments of aphasia prognosis^14,15^ and individual deficits have poor predictive accuracy.^16–18^. Comprehensive models that include characteristics such as stroke location, lesion volume, age, demographics, and initial severity^19–24^ only explain roughly 50% of variance in chronic aphasia severity.^25^

Machine learning (ML) algorithms, which identify patterns in data to make predictions, have been employed to inform aphasia recovery with limited success. These algorithms have incorporated multimodal data including research-grade neuroimaging to augment clinical information^26–28^. However, advanced ML models in aphasia have often used complex structures difficult for users to interpret. They also often rely on data imaging modalities like fMRI that are not widely used in clinical stroke care, which limits their translational potential. A clinically useful algorithm should use readily available data sources and make explainable predictions to enable users to understand the reasoning for guiding decisions.^29,30^ To date, ML development in aphasia has focused on binary classifiers that sort healthy controls from PWA^31,32^ or else broadly group PWA into severe vs mild categories.^28,33,34^ This coarse level of behavioral detail severely limits the ability of these models to inform neurological care and rehabilitation approaches. There remains a need for detailed word-by-word personalized predictions of language deficits in PWA and for these predictions to be both based on clinically accessible data and readily understandable.^29,30^

The linguistic properties of spoken words may be a novel and useful source of multimodal data to predict utterance-specific PWA speech accuracy. Studies in healthy adults and PWA have shown that semantic complexity, the cognitive demands related to selecting between candidate words or recalling specific words, can modulate speech accuracy^35,36^. Metrics related to articulatory difficulty, such as word length in syllables or phonemes, can also be calculated from corpora^37^ of naturalistic spoken English for any word. Incorporating these task-specific linguistic properties^38,39^ along with clinical testing scores, stroke demographics, and neuroimaging, may better capture PWA language performance.^35,38^

Our study aims to predict item-by-item speech ability in chronic post-stroke aphasia, using explainable machine learning on clinically available multimodal input data. We hypothesized that adding linguistic inputs along with PWA clinical details and structural MRI brain networks, would improve word-level predictions of speech accuracy. Using explainability methods, we expected that important features would come from all three sources of multimodal data, including peri-lesional left hemisphere brain regions from canonical language networks, and their intact right hemisphere homotopes. Through web app deployment, we prospectively tested a simplified version of the best model that used only key clinically available inputs. We hypothesized that it would generalize better than the complex classifier for predicting speech of new PWA, both on the same conversational picture description task used to train the model, as well as on different naming tasks using previously unseen English words. Overall, this research integrates multimodal, clinically relevant data to both predict and explain speech deficits, offering a novel aid to patients, families, clinicians, and researchers better understand individual language impairments in people with chronic post-stroke aphasia.

## Methodology

### Retrospective and Prospective Trial Participants

Retrospective data from intake information of a Phase II clinical trial (NCT03651700) included 4620 trials from thirty-five persons with aphasia^40^ (age = 58.7 +/-11, 59% male and 41% female) due to a single left hemisphere stroke, who were in chronic stage (≥6 months post-stroke, lesion overlap seen in Supplemental Figure 1). The data used to test the prospective generalization of the models included 1284 behavioral trials obtained from six PWA recruited later through the same trial, who completed the same language task stimuli, six previously unseen cards, and also a different naming test. All participants had previously been proficient in English and had moderate to severe aphasia as defined by Western Aphasia Battery - Revised Aphasia Quotient (WAB-AQ) scores between 20-85. Present analysis only focuses on neuroimaging and behavioral data from participants’ initial baseline visits. All participants gave informed consent for clinical trial participation, and the study was approved by the Institutional Review Board (831532) at the University of Pennsylvania.

### Language Task and Target Output

Each participant completed a conversational picture description task, with prompts in the style of modified constrain induced language therapy (mCILT)^41^. They were presented with a series of cards depicting common nouns, actions, and scenes and instructed to describe the scene in a sentence to the best of their ability using spoken responses only. This task was geared towards assessing a wide range of everyday conversational and word-finding deficits, differing from traditional picture naming tasks^42,43^ that involve isolated naming of single words from picture cards with less variability in linguistic properties. The language testing cards were balanced for name agreement and noun frequency,^39^ effects that can independently affecting picture naming in PWA. Refer to Supplemental information for details of prompt creation and task evaluation. Resulting PWA utterances (e.g., “the plumber is fixing the pipes”) were recorded and segmented into parts of speech including agent noun, action verb, and object. The noun accuracy was used as the target output measure, i.e., what the machine learning models were trained to predict.

### Multimodal Inputs

Our training and evaluation dataset consisted of 4620 unique trials with 34% correct responses, 66% incorrect responses, and no missing data. Refer to Table 1 for features within the three types of multimodal information: Clinical testing scores and stroke details; Linguistic difficulties from semantic and articulatory corpus data; White matter connectivity strengths between selected brain regions. Refer to Supplemental information for additional details of the data.

**Table 1.** List and Dimensions of Feature Sets before explainability and subset selection.

|  | Subcategory | Feature Descriptions | Dimensions |
| --- | --- | --- | --- |
| **Clinical** | Standardized Language Tests | WAB AQ and WAB sub-scores, Aphasia Type, CCRSA | 6 |
|  | Stroke Information | Lesion volume, time post onset | 2 |
|  | Demographics | Age, gender, solo travel, education, handedness, race, ethnicity | 7 |
| **Linguistics** | Semantic Demands | Word frequency, naming agreement, retrieval, selection | 4 |
|  | Articulatory Demands | Phonemes, morphemes, syllables | 3 |
| **Brain Networks** | DWI MRI based structural connectivity | Strength of each cortical region from Lausanne atlas scale 33 | 83 |

Key: WAB = Western aphasia battery, AQ = aphasia quotient, DWI = diffusion weighted imaging, CCRSA = Communication Confidence Rating Scale for Aphasia

### Linguistic Metrics Calculation from Naturalistic Speech Corpus Data

Lexical frequencies, operationally defined by how often the target word from each picture card occurs in daily language per million spoken words,^44^ were derived from the largest, accuracy-validated English language speech corpus.^37^ Since speech performance in aphasia is known to be influenced by articulatory demands,^45^ we also calculated the number of syllables in a word using the “SyllaPy” package^46,47^ and phonemes using “CMUDict” package^48^, averaged across all acceptable responses. Name agreement, a measure of the proportion of independent subjects providing the same target name as a response to the picture,^49^ was quantified for the agent nouns of each mCILT style language task card only, using norming data from Amazon Mechanical Turk. The number of morphemes for each card were calculated based on consensus ratings from a group including a speech-language pathologist, trained linguist, researchers with experience communicating with PWA, and regional accent speakers (see Supplemental Table 2). Crucially, none of these linguistic metrics are specific to our testing set of agent nouns or cards and can be generated for any new speech prompt.

### White Matter Connectivity Strengths Between Brain Regions

Multi-shell MRI DWI scans were collected within 10 days of language testing for all participants with aphasia (PWA) using a 3T Siemens Prisma FIT VE11C scanner. Refer to Supplemental information for additional MRI protocol details. Note that these scans were structural (not functional), in line with what is typically available in clinical settings. Stroke lesion tracings were created by a neurologist (HBC) and files were then pre-processed to compute tractography, which maps the brain’s white matter pathways using standardized methods (Refer to Supplemental information). Resulting structural brain connectomes were represented as a graph matrix^50^ using volume weighted streamlines based on Lausanne atlas^51^ parcellation at scale 30 (which includes 83 cortical regions). Using commonly accepted network neuroscience methods (Refer to Supplemental information), we computed the weighted strength of each brain region which is a widely characterized measure. A strength of zero implies a structurally disconnected node (such as a cortical region inside a stroke lesion), while greater strength implies a greater structural connectivity of that node.

### Explainable ML Model Development and Evaluation

We trained and optimized machine learning classifiers for supervised binary classification of correct (1) or incorrect (0) speech noun accuracy. Random forest, logistic, regression, support vector machine, and dense neural networks were compared. Model parameters were optimized using standard techniques in a randomized grid search with leave-one-out cross-validation where each machine learning model was trained using all except one random held-out fold for optimization testing and a randomly held out final validation set. Classifier performances was compared for inputs from each of three individual sub-datasets versus in combinations. Refer to Figure 1 for the full machine learning pipeline used. Hypothesis testing was conducted from the model evaluation metrics - Area under the receiver operating curve (AUROC), F1 score, Precision, and Recall - based on 500 iterations of random resampling to ensure replicability.^52^ Statistical comparisons with acceptable type I error for comparing samples^53^ that are normal but not independent were used: Mann-Whitney U test and McNemar’s test for determining whether one learning algorithm outperformed another.^54^ To enhance the interpretability of our best machine learning model’s predictions, we employed a post-hoc explainability algorithm (TreeSHAP)^30^ which uses cooperative game theory^55,56^ to attribute the relative contributions of each feature to the model predictions across all permutations of feature values and combinations. It also provides the directionality of feature’s values with prediction as right or wrong. Feature importances were also compared to a random number feature as a null benchmark.

**Figure 1.**
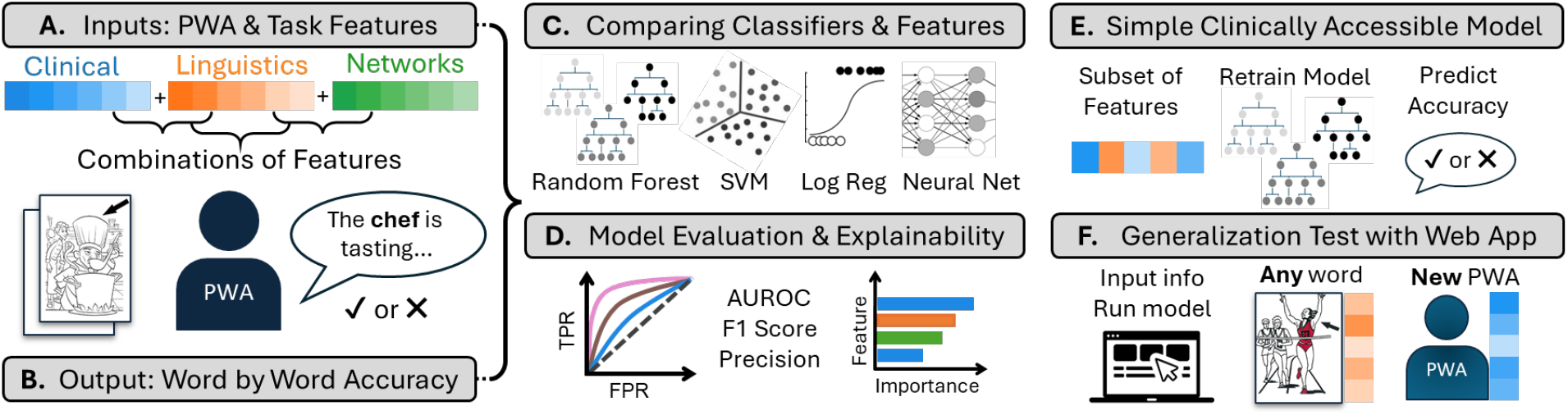
Overview of Methodology. (A) Multimodal inputs consisted of clinical testing scores, linguistic difficulty metrics for each word, and structural network strength of 83 brain regions from each person with aphasia (PWA). (B) Language testing with prompt cards depicting agent nouns performing common actions. Each PWA was shown one card at a time and asked to speak a sentence describing it. Agent noun accuracy on each trial was the target output to be predicted. (C) Overview of ML approach where different classifiers were trained, optimized, and tested using 10 cross validation splits of retrospective data. SVM= support vector machine, Log Reg = logistic regression. (D) Model evaluation metrics AUROC, F1-score, and Precision were computed across 500-fold bootstrapping. Feature importances for the best model were computed using SHAP (Shapley Additive Explanations). (E) A simpler model using clinically available inputs was created and deployed via a web application. (F) Generalization testing using prospective data was conducted for new PWA doing previously unseen prompts as well as entirely new naming tasks.

### Simplified Model and Prospective Testing for Generalization

A simplified version of the best model was created using only widely available input features for ease of clinical applications. It was deployed via a website (https://aphasialens.streamlit.app/) allowing predictions based on user inputs of PWA characteristics and for any word using real-time computation of linguistic metrics from corpus data. Generalization testing set included new PWA who completed picture card description tasks within the same mCILT style for prompts previously unseen in the model training data (new words on same task). Additionally, the model was also tested on a different task (Philadelphia Naming Test). The AUROC scores on prospective generalization were compared to the range of respective models on retrospective testing data from 500 fold bootstrapping distributions.

## Results

### Linguistic Inputs from Naturalistic Speech Corpora Improve Word-by-Word PWA Accuracy Prediction of Random Forest Classifiers

The best models to predict word-level PWA speech accuracies were random forest classifiers using inputs from either all data (mean ± SD for AUROC = 0.867 ± 0.009, F1 score = 0.833 ± 0.071), linguistics plus clinical test scores (AUROC = 0.850 ± 0.066, F1 = 0.795 ± 0.061) or linguistics plus brain network strengths (AUROC = 0.852 ± 0.064, F1 = 0.809 ± 0.063). These models had significantly higher F1 scores than single datasets or combinations without linguistics (Mann Whitey test U statistic range = 4090-6230). Refer to Figure 2. Performance with random forest classifiers was significantly higher (*p*<0.05, McNemar’s test) than support vector classifiers, elastic net logistic regression, and perceptron neural networks as seen in Supplemental Figure 3.

**Figure 2.**
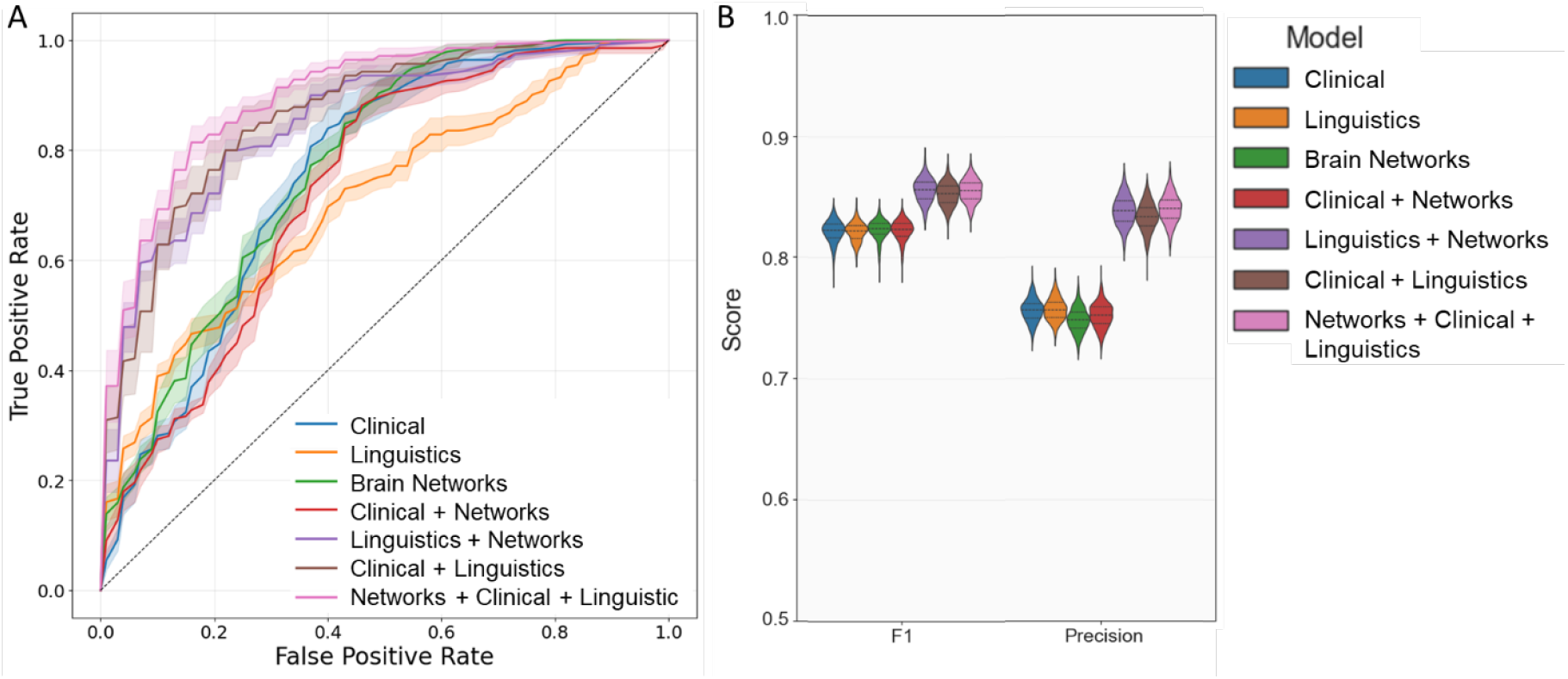
Comparing Models based upon Multimodal Input Features. (A) Clinical Demographics, Linguistic Task Difficulties, Structural Brain Network Strengths, and a combination of the datasets. Receiver Operating Characteristic (ROC) Curves display relative performances compared to chance level as depicted by dashed line. Legend indicates color representations of models. (B) Distributions of bootstrapped model performance metrics.

### Explainable AI Models Highlight Multimodal Feature Importances and Associations with Speech Accuracy Predictions

We used explainable AI to rank feature importances (Refer to Figure 3) and to uncover individual correlations (Supplemental Figure 3) between feature values and influence on model predictions seen through SHAP values. The strongest predictors were WAB-AQ and WAB sub-scores, particularly those related to naming. This was followed by the linguistic priors associated with the semantic demands of the target word: naming agreement and word frequency. Less important linguistic features were number of morphemes, phonemes, and syllables. Key clinical metrics were stroke lesion volume and age, while details such as race, sex, and years of education were not important. Feature explainability of network strengths showed various language network regions highlighted in Figure 3 B.

**Figure 3.**
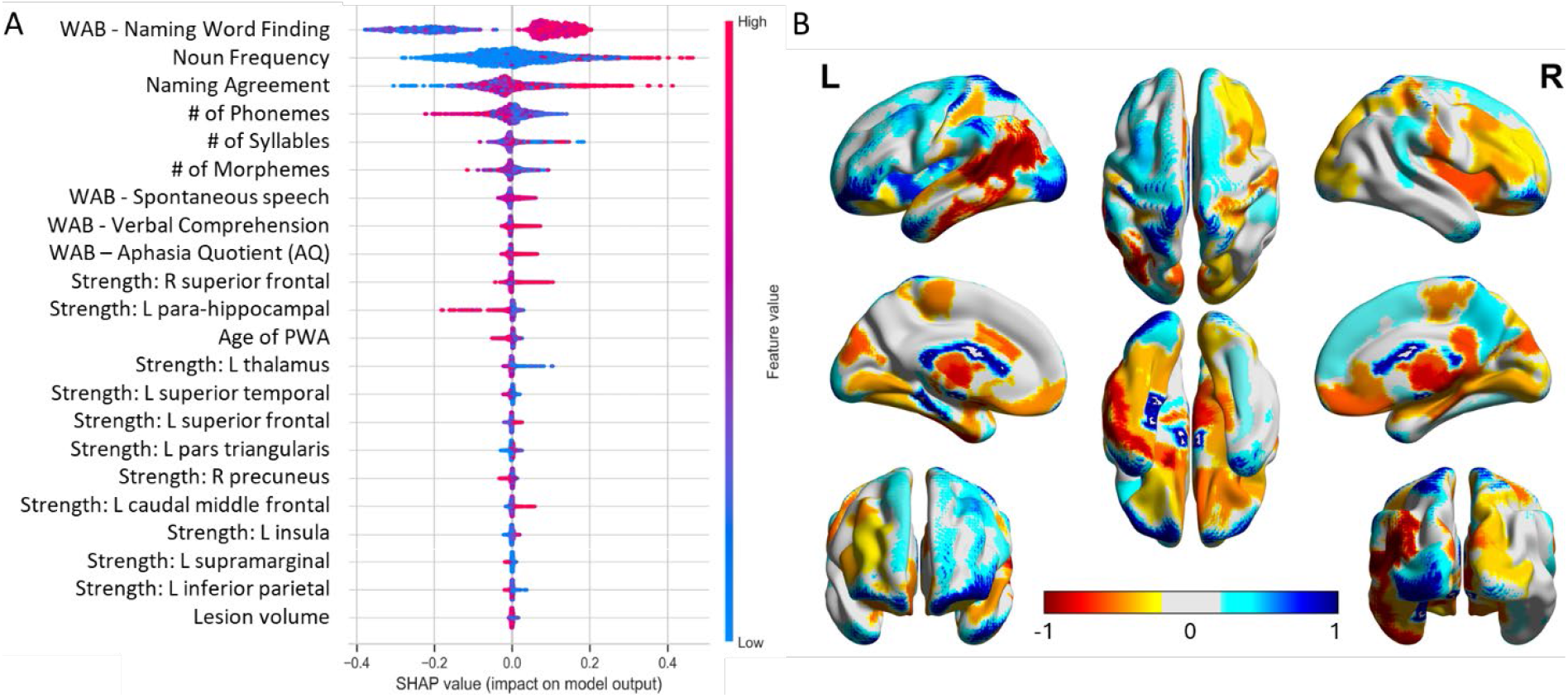
Machine learning model explainability. (A) Feature importances are shown in descending order, with data points colored by feature value distributed across x-axis based upon bootstrapped influence on model predictions towards either correct (positive SHAP values) or incorrect answers (negative SHAP values). (B) Scaled average absolute SHAP values of key regions’ structural connectivity. Warm colors (0.2 to 1.0) indicate strengths associated with greater speech accuracy, while cool colors (-0.2 to -1.0) indicate regions where connectivity is associated with speech deficits.

### Prospective Generalization on New PWA and Unseen Task Prompts

Prospective generalization testing on new PWA with the model using clinical testing and linguistic inputs showed no significant decrease in performance from retrospective testing (*p*>0.05, Mann-Whitney U test). The AUROC mean ± standard error for new PWA doing the same picture description cards was 0.89 ± 0.06 and for the new PWA doing unseen words in the same style of task was 0.87 ± 0.06. Refer to Figure 4A. We found that model performance metrics from generalization test, F1 score, and Precision are significantly (*p*<0.05, *R*^*2*^ >0.4) associated with the difficulty of the card (Refer to Supplemental Figure 4).

**Figure 4.**
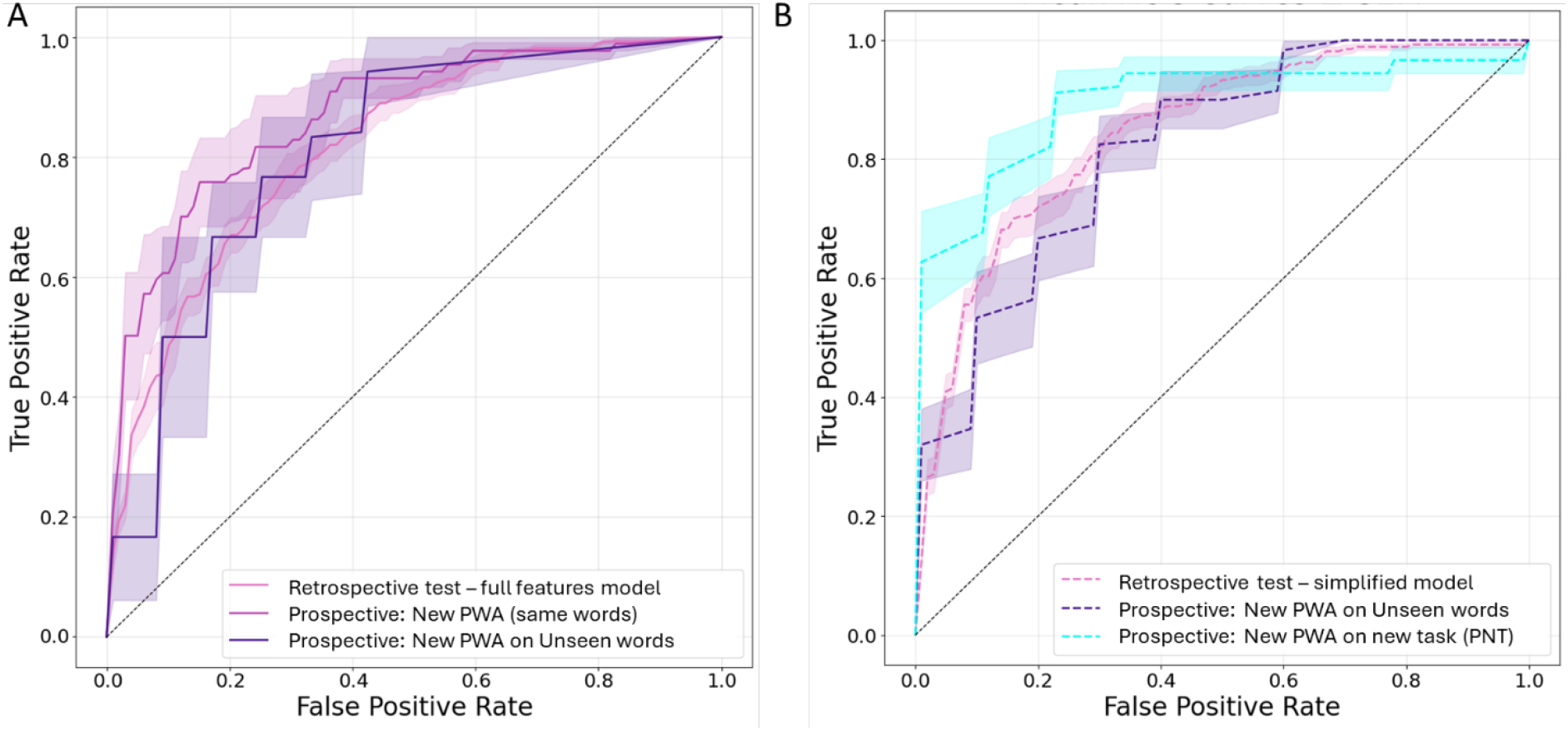
Generalization results with Receiver Operating Characteristic (ROC) Curves. (A) Best full model using all clinical and linguistic information, applied to original testing data (light pink); to new people with aphasia (PWA) doing the same prompts (dark pink); and to new PWA doing previously unseen picture cards (darkest purple). Grey dotted line indicates chance levels. (B) Simplified classifier using a subset of inputs that are widely clinically available, applied on original testing data (light pink dashed); on new PWA doing new cards (dark purple dashed); and on entirely new speech task the Philadelphia Naming Test (light blue dashed line).

### Simplified Model Using Only Clinically Accessible Inputs Predicts PWA Speech Accuracy on Any Word via Web App

Additionally, we created a simpler model using widely clinically available subset of PWA features and linguistic metrics that can be calculated in real time for any user-typed word through a web app (Refer to Supplemental Figure 2 for SHAP importances of simplified features). On retrospective testing, the simpler model had no significant differences in performance (AUROC = 0.84 ± 0.04) than full model (0.86 ± 0.06). On a prospective generalization test to new PWA, the performance was not significantly different from retrospective testing (*p*>0.05, Mann-Whitney U test) with AUROC ± SEM = 0.82 ± 0.07. Additionally, model predictions for a different naming task (Philadelphia Naming Test or PNT) were also not significantly lower (AUROC ± SEM = 0.89 ± 0.08) than for the mCILT style conversational speech test. Refer to Figure 4B.

## Discussion

We created explainable models to predict word-level speech accuracy in chronic post-stroke aphasia with wide generalization to unseen PWA and any new word. This was enabled by combining novel linguistic metrics from corpus data with multimodal neuroimaging and clinical testing features, which to our knowledge surpasses the performance of prior models. Using explainable features to account for the performance of our models makes them more interpretable, which enhances their clinical and translational usefulness. The clinical utility of these results is further aided by our simple, web-run model that requires only a widely available subset of key features for real-time PWA accuracy prediction of naming for any word. Thus, our approach for predicting language performance in PWA is superior in its accuracy, novel in providing item-by-item performance assessments, easily translatable by virtue of using clinically available inputs, applicable to out-of-set information, and transparent in that it provides insights into the factors driving performance classifications.

The clinical implications of predicting word-by-word speech impairments rather than simply assigning labels to individuals are most relevant to guiding neuro-rehabilitation decisions. Recent work has shown that in contrast to historical views of the trajectory of aphasia recovery, chronic aphasia is not static. Additional recovery can occur years after stroke, particularly for persons who receive individualized therapy.^19,26^ However, indexing personalized language deficits in chronic aphasia remains clinically impractical due to limited provider availability and access. While binary predictions of overall aphasia severity - the focus of prior studies - may help to broadly estimate prognosis, they offer limited value for clinicians addressing patients’ day-to-day needs. By contrast, our model can provide insights into the specific communication scenarios where a PWA struggles most, better informing decision-making around priority targets for personalized speech therapy.^57^ In future applications, algorithmic predictions can also be compared to actual performance for tracking changes over time and checking progress in response to speech therapy.

Several of our results extend and complement prior work related to machine learning in aphasia. Prior studies have shown that multimodal data - combining neuroimaging, clinical testing, demographics, and other features - yield the best predictions. ^26,58,59^ Similarly, we show that combining linguistic difficulty with other multimodal inputs yielded better F1 scores than singular datasets. While linguistic metrics were important in all models, there appeared to be redundancy amongst information captured through either clinical testing (lesion volumes and WAB-AQ) or brain network structural connectivity. Recent studies have shown that much of the ‘big data’ benefit from complex neuroimaging can often be explained by simpler lesion volume and location metrics.^60–63^ Likewise, our clinical and network datasets may be capturing duplicate information broadly relating to the extent of the intact language abilities and the damage to language system. We suggest that equivalently performing models using these different inputs can each have unique value for specific applications. For example, explainable models that include detailed structural network features may facilitate the development of novel research hypotheses and testable therapeutic targets related to the neural basis of language and aphasia rehabilitation. Though DWI MRI is growing in clinical availability, model inputs based on standardized clinical testing are likely to have a wider utility and cost- and time-effectiveness. We therefore developed simpler models for prospective generalization testing using a widely available subset of key features from the linguistics and clinical information categories. It can provide real-time insights for clinical decision making in settings where obtaining MRI data might be a limiting factor.^64,65^

Our model explainability results, which include the relationship between the directionality of features with speech abilities, are also highly consistent with prior studies. Underscoring the relevance of standardized batteries, WAB-AQ and its sub-scores were amongst the most important features. Semantic complexity, including word selection and retrieval demands, were most predictive of speech deficits which follows results from healthy adults and PWA responses to therapies.^36,38,40,66–69^ Semantic properties may be a modifiable language attribute for accommodating PWA. In line with prior studies into articulatory difficulty, we also found that phonemes were more predictive of PWA errors than syllable count.^45^ Stroke lesion volume and patient age were also important predictors, especially in the absence of neuroimaging, which is in line with established clinical knowledge.^7,17,18,24^ PWA sex and race did not influence our model predictions which suggests it may have avoided spurious correlations being encoded^70,71^ in line with the mixed results of prior studies.^23,25^ Recent studies of the neural basis of post-stroke function and recovery^61,72^ suggest that not only the lesioned areas, but also intact regions and their network connections with each other are crucial for understanding language and recovery after a stroke.^60,73,74^ We saw the same through explainability analysis: important anatomic regions where greater network strength associated with better speech included left insula and peri-lesional areas. However, right inferior frontal areas homotopic to of lesion were important in the opposite association, wherein greater connectivity predicted more speech deficits. These findings are in line with the inter-hemispheric inhibition hypothesis^75,76^ that suggests a broad network of intact perilesional and contralateral regions are involved in post-stroke aphasia language recovery.^77–79^ The regions associated with worse PWA language may also be candidate sites for inhibitory brain stimulation, which some studies have shown can be a promising aphasia treatment approach.^80,81^

Through our generalization experiment, we addressed a major challenge of machine learning models, which often fail when applied prospectively to out-of-set data.^82,83^ Unlike most studies that train classifiers using retrospective data and test on holdouts, we additionally did a prospective test on new patients recruited at different points in time whose information was preprocessed differently and available only after model completion. We also focused on simplified and explainable AI model to avoid spurious correlations, which have been shown to result in performance over-estimates as high as 20%.^84^ Excellent generalization to new PWA and any word from a naturalistic speech corpus may be partially attributed to our dataset being an order of magnitude larger than any prior ML studies in stroke aphasia in terms of number of unique trials.^26,27,57,85,86^ Other methodologic choices included selecting linguistic metrics which can be generated for most common language prompts through standard open-source software^46–48^ and corpora.^37^ We focused on white matter connectivity from DWI MRI due to its greater clinical availability than fMRI, stability across time^87–90^ and recent inclusion in stroke imaging protocols at high-volume stroke centers.^91–93^ We used stratified cross validation given our mildly imbalanced dataset – a common situation in healthcare applications.^94^ Random forest models do not provide parameter estimates but can be interpretable and explainable post-hoc. This model out-performed other classical machine learning techniques (Supplemental Figure 3) likely due to its robustness to noise and reduced overfitting through ensemble approach.^95^. So for feature selection and explainability we used post-hoc Shapley values^30^ estimation as tree impurity metrics were biased towards higher cardinality features.

We acknowledge limitations related to our study being a secondary analysis of data collected as part of a clinical trial. The PWA research participants may not be fully representative of the chronic post-stroke aphasia population, including those who may have recovered sufficiently or been too impaired to participate in the trial due to additional post-stroke deficits. We focused on agent noun accuracy within the broader communicative utterances as this was most reliable across human scorers. However, future studies may wish to include broader metrics of discourse analysis that can increase external validity. True causal inference remains a goal for the field of machine learning. The Shapley value explanations in this study should thus be interpreted as strong correlations after accounting for co-variates that generate testable hypotheses, not as causal predictors.^65^ Finally, our data were all from chronic PWA, so the model should only be applied and interpreted in this population. Due to greater neuroplasticity in acute stroke, and recovery-related speech changes in subacute phase, we expect altered features to be indicative of speech accuracy in such groups of individuals with aphasia. Future studies should account for temporal inputs into predictions of someone’s aphasia recovery journey such as through large or longitudinal datasets.^94^

This study demonstrates the potential of explainable machine learning in predicting detailed aphasia language abilities, which generalizes to unseen patients and any word using a simple model taking in widely available inputs. This work can help clinical decision making for tailoring speech therapy to patient needs, as well as uncovering research targets to better understand language network and potential neural targets for treatment. It provides a future direction for enhanced precision neurology and efficient stroke rehabilitation, thereby improving outcomes for people with aphasia.

## Data availability

De-identified data of all PWA is available in the Supplemental information section of this paper. Neuroimaging files will be made available only upon request due to privacy restrictions. All code with dependencies related to this study is publicly available at: https://github.com/shreya-parchure/Aphasia-Multimodal-XAI

The web app deployment of the simplified machine learning model associated with this paper is available at: https://aphasialens.streamlit.app/

## Acknowledgements

Authors thank all persons with aphasia who participated and all research, administrative and clinical staff involved with the study.

## References

1. Mukherjee D, Patil CG. Epidemiology and the global burden of stroke. World Neurosurg. 2011;76:S85–90.

2. Tsao CW, Aday AW, Almarzooq ZI, Anderson CAM, Arora P, Avery CL, Baker-Smith CM, Beaton AZ, Boehme AK, Buxton AE, et al. Heart Disease and Stroke Statistics-2023 Update: A Report From the American Heart Association. Circulation. 2023;147:e93–e621.

3. Flowers HL, Skoretz SA, Silver FL, Rochon E, Fang J, Flamand-Roze C, Martino R. Poststroke Aphasia Frequency, Recovery, and Outcomes: A Systematic Review and Meta-Analysis. Arch Phys Med Rehabil. 2016;97:2188-2201.e8.

4. Grönberg A, Henriksson I, Stenman M, Lindgren AG. Incidence of Aphasia in Ischemic Stroke. Neuroepidemiology. 2022;56:174–182.

5. Pedersen PM, Vinter K, Olsen TS. Aphasia after stroke: type, severity and prognosis. The Copenhagen aphasia study. Cerebrovasc Dis. 2004;17:35–43.

6. Hilari K. The impact of stroke: are people with aphasia different to those without? Disabil Rehabil. 2011;33:211–218.

7. Ellis C, Urban S. Age and aphasia: a review of presence, type, recovery and clinical outcomes. Top Stroke Rehabil. 2016;23:430–439.

8. Laska AC, Hellblom A, Murray V, Kahan T, Von Arbin M. Aphasia in acute stroke and relation to outcome. J Intern Med. 2001;249:413–422.

9. Lam JMC, Wodchis WP. The relationship of 60 disease diagnoses and 15 conditions to preference-based health-related quality of life in Ontario hospital-based long-term care residents. Med Care. 2010;48:380–387.

10. Boehme AK, Martin-Schild S, Marshall RS, Lazar RM. Effect of aphasia on acute stroke outcomes. Neurology. 2016;87:2348–2354.

11. Kiran S, Smith ZM. The Efficacy and Utility of Constant Therapy in Poststroke Aphasia Rehabilitation. Perspectives of the ASHA Special Interest Groups. 2025;10:2215–2224.

12. Palmer R, Dimairo M, Cooper C, Enderby P, Brady M, Bowen A, Latimer N, Julious S, Cross E, Alshreef A, et al. Self-managed, computerised speech and language therapy for patients with chronic aphasia post-stroke compared with usual care or attention control (Big CACTUS): a multicentre, single-blinded, randomised controlled trial. Lancet Neurol. 2019;18:821–833.

13. Kristinsson S, den Ouden DB, Rorden C, Newman-Norlund R, Neils-Strunjas J, Fridriksson J. Predictors of Therapy Response in Chronic Aphasia: Building a Foundation for Personalized Aphasia Therapy. J Stroke. 2022;24:189–206.

14. Maas MB, Lev MH, Ay H, Singhal AB, Greer DM, Smith WS, Harris GJ, Halpern EF, Koroshetz WJ, Furie KL. The prognosis for aphasia in stroke. J Stroke Cerebrovasc Dis. 2012;21:350–357.

15. Kertesz A, McCabe P. Recovery patterns and prognosis in aphasia. Brain. 1977;100 Pt 1:1–18.

16. M.M. W, S.A. B. Factors predicting post-stroke aphasia recovery. Journal of the Neurological Sciences. 2015;352:12–18.

17. Thye M, Mirman D. Relative contributions of lesion location and lesion size to predictions of varied language deficits in post-stroke aphasia. Neuroimage Clin. 2018;20:1129–1138.

18. Sperber C, Gallucci L, Mirman D, Arnold M, Umarova RM. Stroke lesion size - Still a useful biomarker for stroke severity and outcome in times of high-dimensional models. Neuroimage Clin. 2023;40:103511.

19. Harvey DY, Parchure S, Hamilton RH. Factors predicting long-term recovery from post-stroke aphasia. Aphasiology. 2022;36:1351–1372.

20. Goldenberg G, Spatt J. Influence of size and site of cerebral lesions on spontaneous recovery of aphasia and on success of language therapy. Brain Lang. 1994;47:684–698.

21. Lazar RM, Minzer B, Antoniello D, Festa JR, Krakauer JW, Marshall RS. Improvement in aphasia scores after stroke is well predicted by initial severity. Stroke. 2010;41:1485–1488.

22. Lazar RM, Speizer AE, Festa JR, Krakauer JW, Marshall RS. Variability in language recovery after first-time stroke. Journal of Neurology, Neurosurgery & Psychiatry. 2008;79:530–534.

23. Plowman E, Hentz B, Ellis C. Post-stroke aphasia prognosis: a review of patient-related and stroke-related factors. Evaluation Clinical Practice. 2012;18:689–694.

24. Benghanem S, Rosso C, Arbizu C, Moulton E, Dormont D, Leger A, Pires C, Samson Y. Aphasia outcome: the interactions between initial severity, lesion size and location. J Neurol. 2019;266:1303–1309.

25. Johnson L, Nemati S, Bonilha L, Rorden C, Busby N, Basilakos A, Newman-Norlund R, Hillis AE, Hickok G, Fridriksson J. Predictors beyond the lesion: Health and demographic factors associated with aphasia severity. Cortex. 2022;154:375–389.

26. Billot A, Lai S, Varkanitsa M, Braun EJ, Rapp B, Parrish TB, Higgins J, Kurani AS, Caplan D, Thompson CK, et al. Multimodal Neural and Behavioral Data Predict Response to Rehabilitation in Chronic Poststroke Aphasia. Stroke. 2022;53:1606–1614.

27. Teghipco A, Newman-Norlund R, Fridriksson J, Rorden C, Bonilha L. Distinct brain morphometry patterns revealed by deep learning improve prediction of post-stroke aphasia severity. Commun Med. 2024;4:1–18.

28. Mahmoud SS, Kumar A, Li Y, Tang Y, Fang Q. Performance Evaluation of Machine Learning Frameworks for Aphasia Assessment. Sensors. 2021;21:2582.

29. Chandler C, Diaz-Asper C, Turner RS, Reynolds B, Elvevåg B. An explainable machine learning model of cognitive decline derived from speech. Alzheimers Dement (Amst). 2023;15:e12516.

30. Lundberg SM, Lee S-I. A Unified Approach to Interpreting Model Predictions [Internet]. In: Guyon I, Luxburg UV, Bengio S, Wallach H, Fergus R, Vishwanathan S, Garnett R, editors. Advances in Neural Information Processing Systems. Curran Associates, Inc.; 2017. Available from: https://proceedings.neurips.cc/paper_files/paper/2017/file/8a20a8621978632d76c43dfd28b67767-Paper.pdf

31. Järvelin A, Juhola M. Comparison of machine learning methods for classifying aphasic and non-aphasic speakers. Computer Methods and Programs in Biomedicine. 2011;104:349–357.

32. Gaspers J, Thiele K, Cimiano P, Foltz A, Stenneken P, Tscherepanow M. An evaluation of measures to dissociate language and communication disorders from healthy controls using machine learning techniques [Internet]. In: Proceedings of the 2nd ACM SIGHIT International Health Informatics Symposium. New York, NY, USA: Association for Computing Machinery; 2012 [cited 2025 Apr 8]. p. 209–218.Available from: https://dl.acm.org/doi/10.1145/2110363.2110389

33. Day M, Dey RK, Baucum M, Paek EJ, Park H, Khojandi A. Predicting Severity in People with Aphasia: A Natural Language Processing and Machine Learning Approach [Internet]. In: 2021 43rd Annual International Conference of the IEEE Engineering in Medicine & Biology Society (EMBC). 2021 [cited 2025 Feb 27]. p. 2299–2302.Available from: https://ieeexplore.ieee.org/abstract/document/9630694?casa_token=-ccXsw4LTRoAAAAA:cUsJ6jrvC63cUOB7S10ghTo7iZGDi2FMN0T9KSu4oztH5soaMmbFGYd_fxqhNVHKOUUE_0A92g

34. Landrigan J-F, Zhang F, Mirman D. A data-driven approach to post-stroke aphasia classification and lesion-based prediction. Brain. 2021;144:1372–1383.

35. McCall J, van der Stelt CM, DeMarco A, Dickens JV, Dvorak E, Lacey E, Snider S, Friedman R, Turkeltaub P. Distinguishing semantic control and phonological control and their role in aphasic deficits: A task switching investigation. Neuropsychologia. 2022;173:108302.

36. Meier EL, Lo M, Kiran S. Understanding semantic and phonological processing deficits in adults with aphasia: Effects of category and typicality. Aphasiology. 2016;30:719–749.

37. Davies M. Word frequency data from The Corpus of Contemporary American English (COCA) [Internet]. 2008 [cited 2025 Apr 7];Available from: https://www.wordfrequency.info/

38. Bastiaanse R, Wieling,Martijn, and Wolthuis N. The role of frequency in the retrieval of nouns and verbs in aphasia. Aphasiology. 2016;30:1221–1239.

39. Almeida J, Knobel M, Finkbeiner M, Caramazza A. The locus of the frequency effect in picture naming: When recognizing is not enough. Psychonomic Bulletin & Review. 2007;14:1177–1182.

40. Dresang HC, Harvey DY, Vnenchak L, Parchure S, Cason S, Twigg P, Faseyitan O, Maher LM, Hamilton RH, Coslett HB. Semantic and Phonological Abilities Inform Efficacy of Transcranial Magnetic Stimulation on Sustained Aphasia Treatment Outcomes. Neurobiol Lang (Camb). 2025;6:nol_a_00160.

41. Maher LM, Kendall D, Swearengin JA, Rodriguez A, Leon SA, Pingel K, Holland A, Rothi LJG. A pilot study of use-dependent learning in the context of Constraint Induced Language Therapy. J Int Neuropsychol Soc. 2006;12:843–852.

42. Lansing AE, Ivnik RJ, Cullum CM, Randolph C. An Empirically Derived Short Form of the Boston Naming Test. Archives of Clinical Neuropsychology. 1999;14:481–487.

43. Walker GM, Schwartz MF. Short-Form Philadelphia Naming Test: Rationale and Empirical Evaluation. American Journal of Speech-Language Pathology. 2012;21:S140–S153.

44. Gertel VH, Karimi H, Dennis NA, Neely KA, Diaz MT. Lexical frequency affects functional activation and accuracy in picture naming among older and younger adults. Psychology and Aging. 2020;35:536–552.

45. Nickels L, and Howard D. Dissociating Effects of Number of Phonemes, Number of Syllables, and Syllabic Complexity on Word Production in Aphasia: It’s the Number of Phonemes that Counts. Cognitive Neuropsychology. 2004;21:57–78.

46. Holtzscher M. mholtzscher/syllapy [Internet]. 2025 [cited 2025 Apr 7];Available from: https://github.com/mholtzscher/syllapy

47. Diaz-Asper M, Holmlund TB, Chandler C, Diaz-Asper C, Foltz PW, Cohen AS, Elvevåg B. Using automated syllable counting to detect missing information in speech transcripts from clinical settings. Psychiatry Res. 2022;315:114712.

48. The CMU Pronouncing Dictionary [Internet]. [cited 2025 Apr 7];Available from: http://www.speech.cs.cmu.edu/cgi-bin/cmudict

49. Barry C, Morrison, C.M., and Ellis AW. Naming the Snodgrass and Vanderwart Pictures: Effects of Age of Acquisition, Frequency, and Name Agreement. The Quarterly Journal of Experimental Psychology Section A. 1997;50:560–585.

50. Bassett DS, Sporns O. Network neuroscience. Nat Neurosci. 2017;20:353–364.

51. Cammoun L, Gigandet X, Meskaldji D, Thiran JP, Sporns O, Do KQ, Maeder P, Meuli R, Hagmann P. Mapping the human connectome at multiple scales with diffusion spectrum MRI. J Neurosci Methods. 2012;203:386–397.

52. Bouckaert RR, Frank E. Evaluating the Replicability of Significance Tests for Comparing Learning Algorithms. In: Dai H, Srikant R, Zhang C, editors. Advances in Knowledge Discovery and Data Mining. Berlin, Heidelberg: Springer; 2004. p. 3–12.

53. Demšar J. Statistical Comparisons of Classifiers over Multiple Data Sets. Journal of Machine Learning Research. 2006;7:1–30.

54. Dietterich TG. Approximate Statistical Tests for Comparing Supervised Classification Learning Algorithms. Neural Computation. 1998;10:1895–1923.

55. Young HP. Monotonic solutions of cooperative games. Int J Game Theory. 1985;14:65–72.

56. Shapley LS. 17. A Value for n-Person Games [Internet]. In: Kuhn HW, Tucker AW, editors. Contributions to the Theory of Games (AM-28), Volume II. Princeton University Press; 1953 [cited 2025 Apr 2]. p. 307–318. Available from: https://www.degruyter.com/document/doi/10.1515/9781400881970-018/html

57. Zhong X. AI-assisted assessment and treatment of aphasia: a review. Front. Public Health [Internet]. 2024 [cited 2025 Apr 8];12. Available from: https://www.frontiersin.org/journals/public-health/articles/10.3389/fpubh.2024.1401240/full

58. White A, Saranti M, d’Avila Garcez A, Hope TMH, Price CJ, Bowman H. Predicting recovery following stroke: Deep learning, multimodal data and feature selection using explainable AI. NeuroImage: Clinical. 2024;43:103638.

59. Olafson ER, Sperber C, Jamison KW, Bowren MD, Boes AD, Andrushko JW, Borich MR, Boyd LA, Cassidy JM, Conforto AB, et al. Data-driven biomarkers better associate with stroke motor outcomes than theory-based biomarkers. Brain Communications. 2024;6:fcae254.

60. Hartwigsen G, Saur D. Neuroimaging of stroke recovery from aphasia - Insights into plasticity of the human language network. Neuroimage. 2019;190:14–31.

61. Yourganov G, Fridriksson J, Rorden C, Gleichgerrcht E, Bonilha L. Multivariate Connectome-Based Symptom Mapping in Post-Stroke Patients: Networks Supporting Language and Speech. J Neurosci. 2016;36:6668–6679.

62. Shewan CM, Kertesz A. Reliability and Validity Characteristics of the Western Aphasia Battery (WAB). Journal of Speech and Hearing Disorders. 1980;45:308–324.

63. Kiran S, Meier EL, Johnson JP. Neuroplasticity in Aphasia: A Proposed Framework of Language Recovery. J Speech Lang Hear Res. 2019;62:3973–3985.

64. Lundberg SM, Nair B, Vavilala MS, Horibe M, Eisses MJ, Adams T, Liston DE, Low DK-W, Newman S-F, Kim J, et al. Explainable machine-learning predictions for the prevention of hypoxaemia during surgery. Nat Biomed Eng. 2018;2:749–760.

65. Ponce-Bobadilla AV, Schmitt V, Maier CS, Mensing S, Stodtmann S. Practical guide to SHAP analysis: Explaining supervised machine learning model predictions in drug development. Clin Transl Sci. 2024;17:e70056.

66. Harvey DY, Traut HJ, Middleton EL. Semantic interference in speech error production in a randomized continuous naming task: Evidence from aphasia. Lang Cogn Neurosci. 2019;34:69–86.

67. Hoffman P, Cogdell-Brooke L, Thompson HE. Going off the rails: Impaired coherence in the speech of patients with semantic control deficits. Neuropsychologia. 2020;146:107516.

68. Bose A, and Schafer G. Name agreement in aphasia. Aphasiology. 2017;31:1143–1165.

69. Kittredge AK, Dell,Gary S., Verkuilen,Jay, and Schwartz MF. Where is the effect of frequency in word production? Insights from aphasic picture-naming errors. Cognitive Neuropsychology. 2008;25:463–492.

70. Huang J, Galal G, Etemadi M, Vaidyanathan M. Evaluation and Mitigation of Racial Bias in Clinical Machine Learning Models: Scoping Review. JMIR Medical Informatics. 2022;10:e36388.

71. Perez-Downes JC, Tseng AS, McConn KA, Elattar SM, Sokumbi O, Sebro RA, Allyse MA, Dangott BJ, Carter RE, Adedinsewo D. Mitigating Bias in Clinical Machine Learning Models. Curr Treat Options Cardio Med. 2024;26:29–45.

72. Koh C-L, Yeh C-H, Liang X, Vidyasagar R, Seitz RJ, Nilsson M, Connelly A, Carey LM. Structural Connectivity Remote From Lesions Correlates With Somatosensory Outcome Poststroke. Stroke. 2021;52:2910–2920.

73. Stockert A, Wawrzyniak M, Klingbeil J, Wrede K, Kümmerer D, Hartwigsen G, Kaller CP, Weiller C, Saur D. Dynamics of language reorganization after left temporo-parietal and frontal stroke. Brain. 2020;143:844–861.

74. Heiss W-D, Thiel A. A proposed regional hierarchy in recovery of post-stroke aphasia. Brain Lang. 2006;98:118–123.

75. van Oers CAMM, Vink M, van Zandvoort MJE, van der Worp HB, de Haan EHF, Kappelle LJ, Ramsey NF, Dijkhuizen RM. Contribution of the left and right inferior frontal gyrus in recovery from aphasia. A functional MRI study in stroke patients with preserved hemodynamic responsiveness. Neuroimage. 2010;49:885–893.

76. van Oers CAMM, van der Worp HB, Kappelle LJ, Raemaekers MAH, Otte WM, Dijkhuizen RM. Etiology of language network changes during recovery of aphasia after stroke. Sci Rep. 2018;8:856.

77. Chu R, Meltzer JA, Bitan T. Interhemispheric interactions during sentence comprehension in patients with aphasia. Cortex. 2018;109:74–91.

78. Turkeltaub PE, Messing S, Norise C, Hamilton RH. Are networks for residual language function and recovery consistent across aphasic patients? Neurology.2011;76:1726–1734.

79. Turkeltaub PE, Coslett HB, Thomas AL, Faseyitan O, Benson J, Norise C, Hamilton RH. The right hemisphere is not unitary in its role in aphasia recovery. Cortex. 2012;48:1179–1186.

80. Hamilton RH, Chrysikou EG, Coslett B. Mechanisms of aphasia recovery after stroke and the role of noninvasive brain stimulation. Brain Lang. 2011;118:40–50.

81. Boddington LJ, Reynolds JNJ. Targeting interhemispheric inhibition with neuromodulation to enhance stroke rehabilitation. Brain Stimulation. 2017;10:214–222.

82. Goetz L, Seedat N, Vandersluis R, van der Schaar M. Generalization—a key challenge for responsible AI in patient-facing clinical applications. npj Digit. Med. 2024;7:1–4.

83. Chekroud AM, Hawrilenko M, Loho H, Bondar J, Gueorguieva R, Hasan A, Kambeitz J, Corlett PR, Koutsouleris N, Krumholz HM, et al. Illusory generalizability of clinical prediction models. Science. 2024;383:164–167.

84. Ong Ly C, Unnikrishnan B, Tadic T, Patel T, Duhamel J, Kandel S, Moayedi Y, Brudno M, Hope A, Ross H, et al. Shortcut learning in medical AI hinders generalization: method for estimating AI model generalization without external data. NPJ Digit Med. 2024;7:124.

85. Pottinger G, Kearns Á. Big data and artificial intelligence in post-stroke aphasia: A mapping review. Advances in Communication and Swallowing. 2024;27:41–55.

86. Kristinsson S, Zhang W, Rorden C, Newman-Norlund R, Basilakos A, Bonilha L, Yourganov G, Xiao F, Hillis A, Fridriksson J. Machine learning-based multimodal prediction of language outcomes in chronic aphasia. Hum Brain Mapp. 2021;42:1682–1698.

87. Abdelnour F, Voss HU, Raj A. Network diffusion accurately models the relationship between structural and functional brain connectivity networks. Neuroimage. 2014;90:335–347.

88. Horn A, Ostwald D, Reisert M, Blankenburg F. The structural-functional connectome and the default mode network of the human brain. Neuroimage. 2014;102 Pt 1:142–151.

89. Hermundstad AM, Bassett DS, Brown KS, Aminoff EM, Clewett D, Freeman S, Frithsen A, Johnson A, Tipper CM, Miller MB, et al. Structural foundations of resting-state and task-based functional connectivity in the human brain. Proc Natl Acad Sci U S A. 2013;110:6169–6174.

90. Lynn CW, Bassett DS. The physics of brain network structure, function and control. Nat Rev Phys. 2019;1:318–332.

91. Hidayat R, Fisher M, Rima SPP, Wiyarta E, Fathi GC, Mustika AP, Irfannadhira AC, Pangeran D, Mesiano T, Kurniawan M, et al. The Necessity of Using MRI as an Imaging Modality in Acute Code Stroke in Indonesia. Vasc Health Risk Manag. 2025;21:207–215.

92. Tan PL, King D, Durkin CJ, Meagher TM, Briley D. Diffusion weighted magnetic resonance imaging for acute stroke: practical and popular. Postgrad Med J. 2006;82:289–292.

93. Schellinger PD, Jansen O, Fiebach JB, Hacke W, Sartor K. A Standardized MRI Stroke Protocol. Stroke. 1999;30:765–768.

94. Tohka J, van Gils M. Evaluation of machine learning algorithms for health and wellness applications: A tutorial. Computers in Biology and Medicine. 2021;132:104324.

95. Breiman L. Random Forests. Machine Learning. 2001;45:5–32.

