## Supplemental information for "Predicting Post-Stroke Aphasia Speech Performance from Multimodal Data with Explainable Machine Learning"

#### Additional Methodology

##### Prompt Creation and Evaluation for mCILT style Language Task

The language task has a constraint to use speech while constructing a full sentence, in the style of modified constraint induced language therapy (mCILT)[1]. To generate the picture cards, we compiled a wordlist of agent nouns that are imageable and had specific labels (i.e., avoiding generic agents such as man/woman in favor of more specific terms like biker/ballerina). Then a Google Image search was completed to locate black and white line drawings of agents clearly engaged in picturable actions that were semantically associated with the agent, consistent with the format seen in other such tasks.[2] Once the images were found, the pictured actions were inventoried in present progressive tense with their associated agent. The task was completed twice, no more than 10 days apart to capture any response inconsistency. Noun accuracy from utterances was scored by blinded, trained raters who reviewed the audio recordings by comparing each spoken noun with the list of acceptable responses for that picture card. Responses were scored as correct if all the phonemes of the target or an acceptable alternative target were produced in the correct order with no erroneously inserted phonemes. Inter-rater and intra-rater reliability were >80% for all PWA transcripts.

##### Structural MRI Acquisition and Processing

Multi-shell MRI DWI scans were collected within 10 days of language testing for all participants with aphasia (PWA) using a 3T Siemens Prisma FIT VE11C scanner. Note that these scans are structural (not functional), in line with what is typically available in clinical settings. Stroke lesion tracings were created by a neurologist (HBC). Preprocessing of the MRI data was conducted on files with imputed lesion area per standard process for stroke lesions.[3] Then QSIPrep 0.13.0RC1, based on Nipype 1.6.0[4] was used to clean the images with MP-PCA denoising, correct B1 field inhomogeneity or distortions, and align the slices. Diffusion images were adjusted to remove noise and standardize the mean intensity of DWI series across separate scanning sequences. Corrections for head motion and eddy currents were performed using FSL,[5] the scan slices were aligned, and outlier replacement was executed (slices with <250 intracerebral voxels or >4 SDs from predictions). Distortion correction was conducted using reversed phase-encode blips, incorporating b=0 reference images. Susceptibility-induced off-resonance fields were estimated as per Smith et al., confounding time-series were calculated, including framewise displacement and head-motion estimates, with DWI resampled to ACPC space (1.3mm isotropic voxels). Thus the final data was aligned to a standard space[6] for subsequent analysis. Tractography reconstruction, which maps the brain's white matter pathways, was conducted in MRtrix3 and using QSIPrep 0.13.0RC1[7] with standard unsupervised algorithms to estimate multi-tissue fiber response functions and fiber orientation directions.[8–10]

#### Computation of Network Connection Strengths

In order to represent structural brain connections as a graph matrix,[11] networks were constructed using volume weighted streamlines for the MRI data based on Lausanne atlas[6] parcellation at scale 30 (which includes 83 cortical regions). This correlation coefficient gives a weighted adjacency matrix $A_{ij}$, wherein $i$ and $j$ are nodes in the matrix connected by edge of weight $a_{ij}$. Nodes within the stroke lesion had all connections set to zero. This network representation fits with common conceptions of the language system as a distributed network within the brain[12,13], and allows for quantitative metrics of neural topology that relate to cognitive properties[14,15] driving complex behaviors like language and recovery from aphasia.[12,16,17]

We computed the strength, a network metric which measures the weighted sum of connections originating from a given node,[18] for all 83 nodes in each person's brain network matrix. Strength of node $i = \Sigma_j a_{ij}$, where $a_{ij}$ represents the weight of the edge connecting node $i$ to node $j$, and the summation is taken across all nodes $j$ connected to node $i$. A strength of zero implies a structurally disconnected node (such as a cortical region inside a stroke lesion), while greater strength implies a greater structural connectivity of that node. Here, connectivity is based on white matter streamlines from a given node with correction for the volume of that cortical region. Strength is a fundamental, well-characterized property of brain networks[19,20] that underlies relationships between structural neuroanatomy and brain dynamics, while also having some correlation with complex network properties like centrality, hub-ness, clustering, modularity, and controllability.[21–24] Thus, we use the 'strength' metric of brain networks as it is a reliable, readily calculated, and understandable measure appropriate for both clinical and basic research. We also created a separate version of the network inputs including only nodes selected by an elastic net algorithm. We created this alternative in case of wide matrix or multicollinearity between the brain network strength value inputs used to train the classifier.

#### Lesion Overlap in Participants

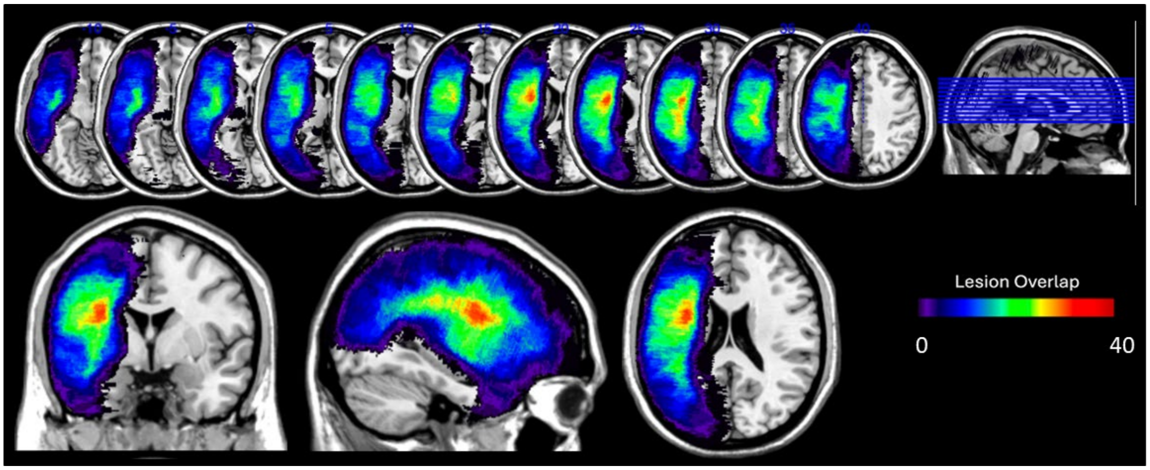


Supplemental Figure 1: Single left hemispheric stroke lesion overlap in participants with aphasia across retrospective and prospective groups.

#### Machine Learning Model Parameter Optimization and Cross-validation

After selecting parameters with random grid search on nested cross-validation we evaluated only the best models of each type. We used stratified *k*-fold cross-validation, a resampling procedure where the training set was randomly split without resampling into *k*- smaller sets having the same stratification rate of each class and independently evaluated on all. This is considered the gold standard method for model evaluation especially to deal with imbalanced data as was the case in our set and performs better than holdout.[25] All folds of the dataset were stratified to ensure the same distribution of correct and incorrect answers. Random forests are an ensemble learning technique that classifies data by constructing independent decision trees each trained on a bootstrap sample of the data and aggregating their predictions to make a final classification.[26] This model was chosen as it handles multimodal data inputs with differing scales and cardinalities, has inbuilt bagging and bootstrapping to prevent overfitting, and is robust to noise even with smaller datasets.

#### Machine Learning Model Evaluation Metrics

The optimized models were evaluated on held-out validation data (random 10% of the 4620 trials stratified to have the same base rate). We compared models' abilities to distinguish between classes at a range of detection thresholds using Area Under the Curve (AUC). Precision was used to indicate the proportion of true positive predictions among all positive predictions, while Recall reflected the model's ability to identify all relevant instances – both computed for the incorrect answers class. We also computed the F1 score, which balances precision and recall, in addition to Accuracy weighted by class.[25] Further, bootstrapped performance was computed across 1000 5-fold random cross validation splits of the data using the best model to train and test independently in each partition.[27] From this, a distribution of values for each metric was obtained.

#### Feature Importances and AI model Explainability

To enhance the interpretability of our best machine learning model's predictions, we employed a post-hoc explainability algorithm (SHAP) based upon Shapley values,[28] which has had recent success in healthcare AI applications[29]. Shapley values are derived from cooperative game theory,[30,31] and here attribute the contribution of each feature to the model's output based on its marginal contributions across all possible combinations of features. This method provides a nuanced understanding of feature interactions and their individual impacts on predictions, facilitating deeper insights into the model's decision-making process. Evaluating feature importance using permutations testing and game theory-based Shapley values of causal contribution to model predictions shows not just that a feature is important, but also the direction of its relationship with respect to predicted improving or worsening language impairments. Shapley values track with permutation importance based on how dropping a feature impacts model accuracy, while also additionally sampling the full range of feature values and how that shifts model predictions. Feature importances were also compared to a random number feature as null model benchmark.

#### Simplified Model and Web Application Deployment

To create a widely usable model that will make predictions for any user-input word, we used simplified input features based on information that would be readily available in a clinical setting. These include 9 features where 6 are PWA information to be entered and 3 are linguistic metrics calculated in real time for the typed word.

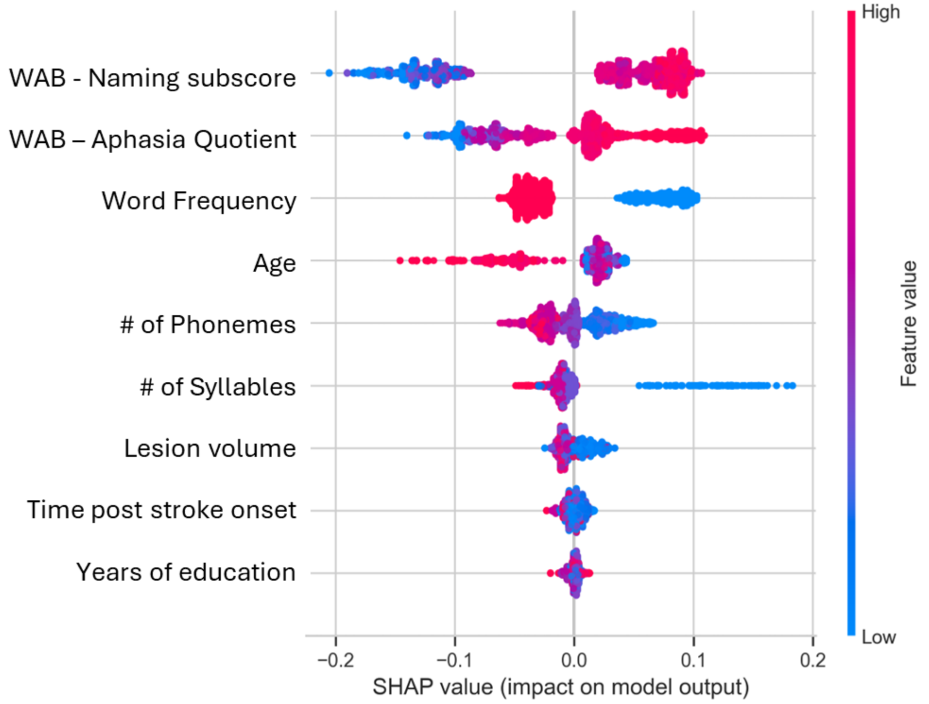


Supplemental Figure 2. Simplified Classifier Using Subset of Inputs that are widely Clinically available. Feature importances are in descending order based on mean absolute SHAP values.

### Supplemental Results

#### Random forests outperform other model types

Using all input features to train various AI model types with parameter optimization, we found that trial-level accuracy of agent noun communication exchange in chronic aphasia was best predicted by the Random Forest classifier (AUC ± cross validation SEM = 0.89 ± 0.06) and Logistic Regression (0.89 ± 0.06) followed by the Neural Network (0.85 ± 0.06), Support Vector Machine (0.82 ± 0.06), and Elastic Net (0.82 ± 0.04) model types.

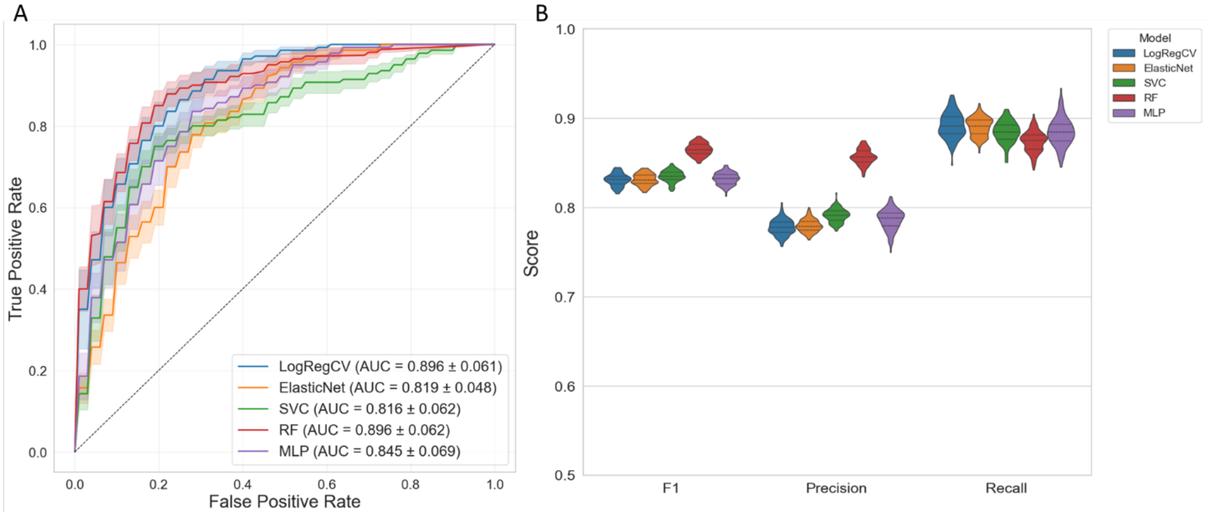


Supplemental Figure 3: Comparing different types of ML algorithms. (A) ROC curves with standard deviations across ten stratified cross validation splits. (B) Distributions of 500 fold bootstrapped performance metrics for optimized model. Key: LogRegCV = logistic regression; SVC = support vector classifier; RF = random forest; MLP = multi-layer perceptron.

#### Association between word difficulty and generalization performance

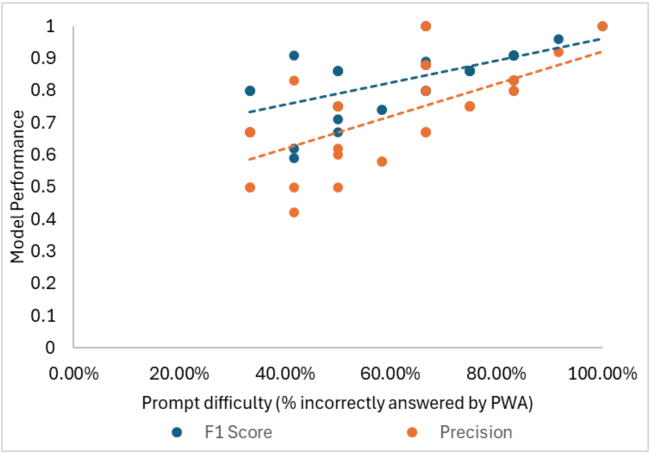


Supplemental Figure 4. Generalization test metrics F1 score and Precision were significantly ($p$<0.05, $R^2$ >0.4) associated with the difficulty of the card.

### Supplemental Tables

##### Supplemental Table 1. Participant demographics & aphasia characteristics

| Subject ID | MPO | Age | Sex | WAB AQ | CCRSA | Aphasia Type | Lesion Volume (cc) |
|---|---|---|---|---|---|---|---|
| TMSA-02 | 100 | 55 | Male | 77 | 22 | Broca's | 100500 |
| TMSA-04* | 45 | 75 | Male | 31 | 24 | Broca’s | 106500 |
| TMSA-06 | 20 | 60 | Male | 84 | 32 | Conduction | 162000 |
| TMSA-11 | 30 | 55 | Male | 34 | 37 | Broca's | 190200 |
| TMSA-13* | 100 | 60 | Male | 85 | 26 | Anomic | 26000 |
| TMSA-15 | 110 | 60 | Female | 76 | 18 | Broca's | 52500 |
| TMSA-19 | 240 | 75 | Male | 53 | 34 | Broca's | 285600 |
| TMSA-20 | 50 | 70 | Female | 70 | 35 | Conduction | 51700 |
| TMSA-25 | 180 | 65 | Male | 77 | 32 | Conduction | 181100 |
| TMSA-24* | 20 | 60 | Male | 20 | 26 | Broca’s | 124000 |
| TMSA-26* | 130 | 75 | Male | 45 | 36 | Wernicke's | 109000 |
| TMSA-27 | 80 | 55 | Male | 40 | 16 | Broca's | 89600 |
| TMSA-28 | 10 | 55 | Male | 65 | 14 | Conduction | 18600 |
| TMSA-33 | 20 | 50 | Male | 49 | 25 | Broca's | 175900 |
| TMSA-35 | 160 | 60 | Male | 84 | 38 | Anomic | 38700 |
| TMSA-36 | 20 | 65 | Male | 80 | 25 | Conduction | 44600 |
| TMSA-38 | 10 | 70 | Male | 80 | 16 | Anomic | 152200 |
| TMSA-39 | 20 | 60 | Female | 77 | 33 | Broca's | 24100 |
| TMSA-40 | 10 | 65 | Male | 83 | 23 | Anomic | 74500 |
| TMSA-43 | 20 | 70 | Male | 73 | 25 | Conduction | 23500 |
| TMSA-44 | 30 | 70 | Male | 81 | 25 | Anomic | 79400 |
| TMSA-47 | 10 | 65 | Female | 76 | 15 | Anomic | 2200 |
| TMSA-48 | 10 | 65 | Male | 65 | 22 | Broca's | 202500 |
| TMSA-52 | 10 | 60 | Female | 80 | 21 | Anomic | 67100 |
| TMSA-54 | 50 | 35 | Male | 63 | 24 | Anomic | 230800 |
| TMSA-55 | 40 | 70 | Male | 50 | 32 | Transcortical Sensory | 106900 |
| TMSA-58 | 20 | 75 | Male | 63 | 29 | Broca's | 17800 |
| TMSA-59* | 20 | 50 | Female | 83 | 27 | Anomic | 54600 |
| TMSA-63 | 180 | 35 | Female | 75 | 28 | Conduction | 215400 |
| TMSA-64 | 90 | 35 | Male | 84 | 35 | Anomic | 27100 |
| TMSA-66 | 50 | 50 | Female | 85 | 34 | Anomic | 21100 |
| TMSA-67 | 40 | 60 | Female | 84 | 30 | Anomic | 28700 |
| TMSA-73 | 40 | 60 | Female | 84 | 16 | Anomic | 119800 |
| TMSA-74 | 290 | 60 | Female | 80 | 39 | Transcortical Motor | 64200 |
| TMSA-78 | 10 | 50 | Female | 72 | 31 | Conduction | 21100 |
| TMSA-79 | 80 | 75 | Female | 77 | 12 | Transcortical Motor | 67300 |
| TMSA-80 | 30 | 70 | Male | 71 | 40 | Wernicke's | 32900 |
| TMSA-81 | 70 | 65 | Female | 81 | 36 | Anomic | 30600 |
| TMSA-84 | 10 | 45 | Female | 78 | 32 | Conduction | 10300 |
| TMSA-86* | 240 | 55 | Female | 75 | 30 | Broca's | 101300 |

Note: MPO = months post onset of stroke, WAB AQ = Western Aphasia Battery Aphasia Quotient, CCRSA = Communication Confidence Rating Scale for Aphasia. All numerical values are rounded to preserve anonymity. Subject ID followed by * indicates participants who were part of prospective generalization.

**Supplemental Table 2. Linguistic priors for each communicative exchange agent noun**

| Agent Noun | Word Frequency | Naming Agreement | Syllables | Phonemes | Morphemes |
|---|---|---|---|---|---|
| anchorwoman | LF | LA | 4 | 9 | 2 |
| archeologist | LF | HA | 5 | 11 | 2.5 |
| architect | LF | HA | 3 | 8 | 2 |
| artist | HF | LA | 2 | 6 | 2 |
| astronaut | HF | HA | 3 | 8 | 2 |
| ballerina | LF | HA | 4 | 7 | 2 |
| bank teller | LF | HA | 3 | 8 | 2.5 |
| barber | HF | HA | 2 | 5 | 1.5 |
| bartender | HF | HA | 3 | 8 | 3 |
| bellhop | LF | LA | 2 | 6 | 2 |
| blacksmith | LF | HA | 2 | 8 | 2 |
| boyscout | LF | LA | 2 | 6 | 2 |
| butcher | HF | HA | 2 | 4 | 1.5 |
| cameraman | LF | HA | 4 | 8 | 2 |
| captain | HF | LA | 2 | 6 | 1 |
| carpenter | LF | LA | 3 | 8 | 1.5 |
| caveman | LF | HA | 2 | 6 | 2 |
| chef | HF | HA | 1 | 3 | 1 |
| chimney sweep | LF | HA | 3 | 9 | 2 |
| construction worker | LF | LA | 5 | 15 | 4 |
| cyclist | LF | HA | 2.5 | 7 | 2 |
| dentist | HF | HA | 2 | 7 | 2 |
| detective | HF | HA | 3 | 8 | 2 |
| doctor | HF | HA | 2 | 5 | 2 |
| explorer | HF | LA | 3 | 8 | 2 |
| farmer | HF | HA | 2 | 5 | 2 |
| figure skater | LF | HA | 4 | 10 | 3 |
| fisherman | HF | HA | 3 | 7 | 2.5 |
| garbageman | LF | LA | 3 | 9 | 2 |
| gardener | LF | HA | 3 | 7 | 2 |
| goalie | LF | HA | 2 | 4 | 2 |
| golfer | HF | HA | 2 | 5 | 2 |
| guitarist | HF | LA | 3 | 8 | 2 |
| hairdresser | LF | LA | 3 | 8 | 3 |
| hunter | HF | HA | 2 | 5 | 2 |
| janitor | LF | LA | 3 | 6 | 2 |
| jester | LF | LA | 2 | 5 | 2 |
| jockey | HF | HA | 2 | 4 | 1 |
| judge | HF | HA | 1 | 3 | 1 |
| jury | HF | HA | 2 | 4 | 1 |
| knight | LF | LA | 1 | 3 | 1 |
| landscaper | LF | LA | 3 | 9 | 3 |
| lineman | LF | LA | 2 | 6 | 2 |
| mailman | LF | LA | 2 | 6 | 2 |
| mason | LF | LA | 2 | 5 | 2 |
| mechanic | HF | HA | 3 | 7 | 1.5 |
| mover | LF | HA | 2 | 4 | 2 |
| nun | HF | HA | 1 | 3 | 1 |
| pharmacist | HF | HA | 3 | 9 | 3 |
| photographer | HF | HA | 4 | 9 | 3 |
| pitcher | HF | HA | 2 | 4 | 2 |
| police | HF | HA | 2 | 5 | 1 |
| potter | LF | HA | 2 | 4 | 2 |

| | | | | | |
|---|---|---|---|---|---|
| priest | HF | HA | 2 | 5 | 1.5 |
| receptionist | LF | LA | 4 | 11 | 2 |
| runner | HF | LA | 2 | 4 | 2 |
| scientist | HF | HA | 3 | 8 | 2 |
| sculptor | LF | HA | 2 | 7 | 2 |
| skydiver | LF | LA | 3 | 7 | 3 |
| student | HF | HA | 2 | 7 | 2 |
| tailor | LF | HA | 2 | 4 | 1.5 |
| thief | HF | HA | 1.5 | 3 | 1.5 |
| veterinarian | LF | HA | 4 | 11 | 1.5 |
| waitress | HF | HA | 2 | 6 | 2 |
| witch | HF | HA | 1 | 3 | 1 |
| witness | HF | HA | 2 | 6 | 1 |

Note: HF = high frequency, HA = high agreement, LF = low frequency, LA = low agreement. Syllables, Phonemes, and Morphemes were averaged based on the different target responses accepted and differences in pronunciation.

### Supplemental References


1. Maher LM, Kendall D, Swearengin JA, Rodriguez A, Leon SA, Pingel K, Holland A, Rothi LJG. A pilot study of use-dependent learning in the context of Constraint Induced Language Therapy. *J Int Neuropsychol Soc*. 2006;12:843–852.

2. Edmonds LA, Nadeau SE, Kiran S. Effect of Verb Network Strengthening Treatment (VNeST) on Lexical Retrieval of Content Words in Sentences in Persons with Aphasia. *Aphasiology*. 2009;23:402–424.

3. Erickson BA, Kim B, Deck BL, Pustina D, DeMarco AT, Dickens JV, Kelkar AS, Turkeltaub PE, Medaglia JD. Preserved anatomical bypasses predict variance in language functions after stroke. *Cortex*. 2022;155:46–61.

4. Gorgolewski K, Burns CD, Madison C, Clark D, Halchenko YO, Waskom ML, Ghosh SS. Nipype: a flexible, lightweight and extensible neuroimaging data processing framework in python. *Front Neuroinform*. 2011;5:13.

5. Jenkinson M, Beckmann CF, Behrens TEJ, Woolrich MW, Smith SM. FSL. *NeuroImage*. 2012;62:782–790.

6. Cammoun L, Gigandet X, Meskaldji D, Thiran JP, Sporns O, Do KQ, Maeder P, Meuli R, Hagmann P. Mapping the human connectome at multiple scales with diffusion spectrum MRI. *J Neurosci Methods*. 2012;203:386–397.

7. Cieslak M, Cook PA, He X, Yeh F-C, Dhollander T, Adebimpe A, Aguirre GK, Bassett DS, Betzel RF, Bourque J, et al. QSIPrep: an integrative platform for preprocessing and reconstructing diffusion MRI data. *Nat Methods*. 2021;18:775–778.

8. Tournier J-D, Calamante F, Gadian DG, Connelly A. Direct estimation of the fiber orientation density function from diffusion-weighted MRI data using spherical deconvolution. *Neuroimage*. 2004;23:1176–1185.

9. Tournier J-D, Calamante F, Connelly A. Robust determination of the fibre orientation distribution in diffusion MRI: non-negativity constrained super-resolved spherical deconvolution. *Neuroimage*. 2007;35:1459–1472.

10. Dhollander T, Clemente A, Singh M, Boonstra F, Civier O, Duque JD, Egorova N, Enticott P, Fuelscher I, Gajamange S, et al. Fixel-based Analysis of Diffusion MRI: Methods, Applications, Challenges and Opportunities. *NeuroImage*. 2021;241:118417.

11. Bassett DS, Sporns O. Network neuroscience. *Nat Neurosci*. 2017;20:353–364.

12. Yourganov G, Fridriksson J, Rorden C, Gleichgerrcht E, Bonilha L. Multivariate Connectome-Based Symptom Mapping in Post-Stroke Patients: Networks Supporting Language and Speech. *J Neurosci*. 2016;36:6668–6679.

13. Fedorenko E, Ivanova AA, Regev TI. The language network as a natural kind within the broader landscape of the human brain. *Nat. Rev. Neurosci.* 2024;25:289–312.

14. Bassett DS, Mattar MG. A Network Neuroscience of Human Learning: Potential To Inform Quantitative Theories of Brain and Behavior. *Trends Cogn Sci*. 2017;21:250–264.

15. Medaglia JD, Pasqualetti F, Hamilton RH, Thompson-Schill SL, Bassett DS. Brain and cognitive reserve: Translation via network control theory. *Neurosci Biobehav Rev*. 2017;75:53–64.

16. Hartwigsen G, Saur D. Neuroimaging of stroke recovery from aphasia - Insights into plasticity of the human language network. *Neuroimage*. 2019;190:14–31.

17. Kiran S, Thompson CK. Neuroplasticity of Language Networks in Aphasia: Advances, Updates, and Future Challenges. *Front Neurol*. 2019;10:295.

18. Gusfield D. Computing the Strength of a Graph. *SIAM J. Comput.* 1991;20:639–654.

19. Sporns O. Graph theory methods: applications in brain networks. *Dialogues in Clinical Neuroscience*. 2018;20:111–121.

20. Sporns O, Tononi G, Edelman GM. Connectivity and complexity: the relationship between neuroanatomy and brain dynamics. *Neural Networks*. 2000;13:909–922.

21. Rubinov M, Sporns O. Complex network measures of brain connectivity: Uses and interpretations. *NeuroImage*. 2010;52:1059–1069.

22. Medaglia JD, Harvey DY, Kelkar AS, Zimmerman JP, Mass JA, Bassett DS, Hamilton RH. Language Tasks and the Network Control Role of the Left Inferior Frontal Gyrus. *eNeuro*. 2021;8:ENEURO.0382-20.2021.

23. Medaglia JD, Harvey DY, White N, Kelkar A, Zimmerman J, Bassett DS, Hamilton RH. Network Controllability in the Inferior Frontal Gyrus Relates to Controlled Language Variability and Susceptibility to TMS. *J. Neurosci.* 2018;38:6399–6410.

24. Sporns O, Honey CJ, Kötter R. Identification and Classification of Hubs in Brain Networks. *PLOS ONE*. 2007;2:e1049.

25. Tohka J, van Gils M. Evaluation of machine learning algorithms for health and wellness applications: A tutorial. *Computers in Biology and Medicine*. 2021;132:104324.

26. Breiman L. Random Forests. *Machine Learning*. 2001;45:5–32.

27. Rainio O, Teuho J, Klén R. Evaluation metrics and statistical tests for machine learning. *Sci Rep*. 2024;14:6086.

28. Lundberg SM, Lee S-I. A Unified Approach to Interpreting Model Predictions [Internet]. In: Guyon I, Luxburg UV, Bengio S, Wallach H, Fergus R, Vishwanathan S, Garnett R, editors. Advances in Neural Information Processing Systems. Curran Associates, Inc.; 2017. Available from: https://proceedings.neurips.cc/paper_files/paper/2017/file/8a20a8621978632d76c43dfd28b67767-Paper.pdf

29. Ponce-Bobadilla AV, Schmitt V, Maier CS, Mensing S, Stodtmann S. Practical guide to SHAP analysis: Explaining supervised machine learning model predictions in drug development. *Clin Transl Sci*. 2024;17:e70056.

30. Young HP. Monotonic solutions of cooperative games. *Int J Game Theory*. 1985;14:65–72.

31. Shapley LS. 17. A Value for n-Person Games [Internet]. In: Kuhn HW, Tucker AW, editors. Contributions to the Theory of Games (AM-28), Volume II. Princeton University Press; 1953 [cited 2025 Apr 2]. p. 307–318.Available from: https://www.degruyter.com/document/doi/10.1515/9781400881970-018/html
